# Long read sequencing reveals Poxvirus evolution through rapid homogenization of gene arrays

**DOI:** 10.1101/245373

**Authors:** Thomas A. Sasani, Kelsey R. Cone, Aaron R. Quinlan, Nels C. Elde

## Abstract

Large DNA viruses rapidly evolve to defeat host defenses. Poxvirus adaptation can involve combinations of recombination-driven gene copy number variation and beneficial single nucleotide variants (SNVs) at the same locus, yet how these distinct mechanisms of genetic diversification might simultaneously facilitate adaptation to immune blocks is unknown. We performed experimental evolution with a vaccinia virus population harboring a SNV in a gene actively undergoing copy number amplification. Comparisons of virus genomes using the Oxford Nanopore Technologies sequencing platform allowed us to phase SNVs within large gene copy arrays for the first time, and uncovered a mechanism of adaptive SNV homogenization reminiscent of gene conversion, which is actively driven by selection. Our work reveals a new mechanism for the fluid gain of beneficial mutations in genetic regions undergoing active recombination in viruses, and illustrates the value of long read sequencing technologies for investigating complex genome dynamics in diverse biological systems.

## Introduction

Gene duplication is long recognized as a potential source of genetic innovation (Ohno 1970). Following duplication events, the resulting stretches of homologous sequence can promote recombination between gene copies. Gene conversion, the nonreciprocal transfer of sequence between homologous genetic regions, is one outcome of recombination evident in diverse eukaryotes (Brown et al., 1972; Semple and Wolfe, 1999; Drouin, 2002; Rozen et al., 2003; Ezawa et al., 2006; Chen et al., 2007), as well as in bacterial and archaeal genomes (Santoyo and Romero, 2005; Soppa, 2011). Gene conversion can result in high homology among duplicated gene copies, as in ribosomal RNA gene arrays (Liao, 1999; Eickbush and Eickbush, 2007). However, in other cases, such as the human leukocyte antigen gene family (Zangenberg et al., 1995) or the transmembrane protein gene cassettes of some pathogenic bacteria (Santoyo and Romero, 2005), gene conversion can also generate sequence diversity. Because multiple gene copies create more targets for mutation, variants that arise within individual copies can be efficiently spread or eliminated through gene conversion (Mano and Innan, 2008; Ellison and Bachtrog, 2015).

Although recombination might influence genetic variation in populations on very short time scales, many studies to date have used phylogenetic analysis in relatively slow-evolving populations to infer outcomes of gene conversion, including cases of concerted evolution, where homology of gene families within a species exceeds the homology of orthologous genes between species (Chen et al., 2007; Ohta, 2010). Extensive studies in yeast and bacteria (reviewed in Petes and Hill, 1988; Perkins, 1992; Haber, 2000; Santoyo and Romero, 2005), and some recent work in viruses (Hughes, 2004; Ba abdullah et al., 2017), consider gene conversion on shorter time scales. In order to expand our understanding of how recombination might influence virus variation during the course of adaptation, we focused on large DNA viruses, in which rapidly evolving populations can simultaneously harbor both adaptive gene copy number variation and beneficial single nucleotide variants (SNVs) at the same locus.

Poxviruses are an intriguing system to study mechanisms of rapid adaptation, as they possess high rates of recombination (Ball, 1987; Evans et al., 1988; Spyropoulos et al., 1988; Merchlinsky, 1989) that lead to the recurrent emergence of tandem gene duplications (Slabaugh et al., 1989; Elde et al., 2012; Brennan et al., 2014; Erlandson et al., 2014; Cone et al., 2017;). The poxvirus DNA polymerase gene encodes both replicase and recombinase activities, reflecting a tight coupling of these essential functions for virus replication (Colinas et al., 1990; Willer et al., 1999; Hamilton and Evans, 2005). Polymerase-associated recombination may underlie the rapid appearance of gene copy number variation (CNV), which was proposed as a potentially widespread mechanism of vaccinia virus adaptation following courses of experimental evolution (Elde et al., 2012, Cone et al., 2017). In these studies, recurrent duplications of the K3L gene, which encodes a weak inhibitor of the host innate immune factor Protein Kinase R (PKR; Davies et al., 1992), were identified following serial infections of human cells with a vaccinia strain lacking a strong PKR inhibitor encoded by the E3L gene (ΔE3L; Chang et al., 1992; Beattie et al., 1995). In addition to copy number amplification, a beneficial single nucleotide variant arose in the K3L gene in some populations, resulting in a His47Arg amino acid change (K3L^His47Arg^), which encodes enhanced inhibition of PKR activity and aids virus replication (Kawagishi-Kobayashi et al., 1997; Elde et al., 2012). However, the mechanisms by which these distinct mechanisms of adaptation might synergize or compete in virus populations during virus evolution are unknown.

To investigate how heterogeneous virus populations adapt to cellular defenses, we performed courses of experimental evolution with a vaccinia virus population containing both K3L CNV and the K3L^His47Arg^ SNV (Elde et al., 2012). To overcome the challenge of genotyping point mutations in repetitive arrays of the K3L gene, we sequenced virus genomes with the Oxford Nanopore Technologies (ONT) MinION platform. The MinION platform, which is capable of sequencing extremely long DNA molecules (Jain et al., 2017), allowed us to develop an integrated pipeline to analyze CNV at single-genome scale. Long sequencing reads, which can completely span tandem arrays of K3L duplications (up to 15 copies, and 99 kbp in this study), allowed us to track the K3L^His47Arg^ mutation as it spread through K3L gene arrays within an evolving virus population. Altering conditions in this experimental system allowed us to assess the impact of selection and recombination on K3L^His47Arg^ variant accumulation within K3L arrays. These analyses of variant dynamics reveal a viral mechanism of genetic homogenization resembling gene conversion, and demonstrate how long read sequencing can facilitate studies of recombination-driven genome evolution.

## Results

### Accumulation of a single nucleotide variant during gene copy number variation

In previous work, we collected a virus population adapted over ten serial infections that contains gene copy number amplifications of K3L, and the beneficial K3L^His47Arg^ point mutation at roughly 10% frequency in the population (**Figure 1A, 1B**, up to passage 10; Elde et al., 2012). To study the fate of the K3L^His47Arg^ variant among repetitive arrays of K3L, we performed ten additional serial infection passages (P11-P20) in human cells. Virus fitness, as judged by comparative replication in human cells, remained well above parent (ΔE3L) levels through P20 (**Figure 1A**), consistent with an ongoing course of virus adaptation involving variation in K3L. The emergence of K3L CNV within virus genomes coincided with an increase in fitness around P5 (Elde et al., 2012), and gene copy number increases appeared to stabilize by P10 (**Figure 1B**). Notably, the K3L^His47Arg^ SNV, though apparently stable at an estimated frequency of 0.1 in the population from P5 through P10 (Elde et al., 2012), accumulated to near fixation between P10 and P20 (frequency of approximately 0.9; **Figure 1B**). Thus, despite a plateau in fitness across later experimental passages, the accumulation of the beneficial point mutation suggests that selection still acts on the K3L^His47Arg^ variant in the heterogeneous virus population.

**Figure 1.**
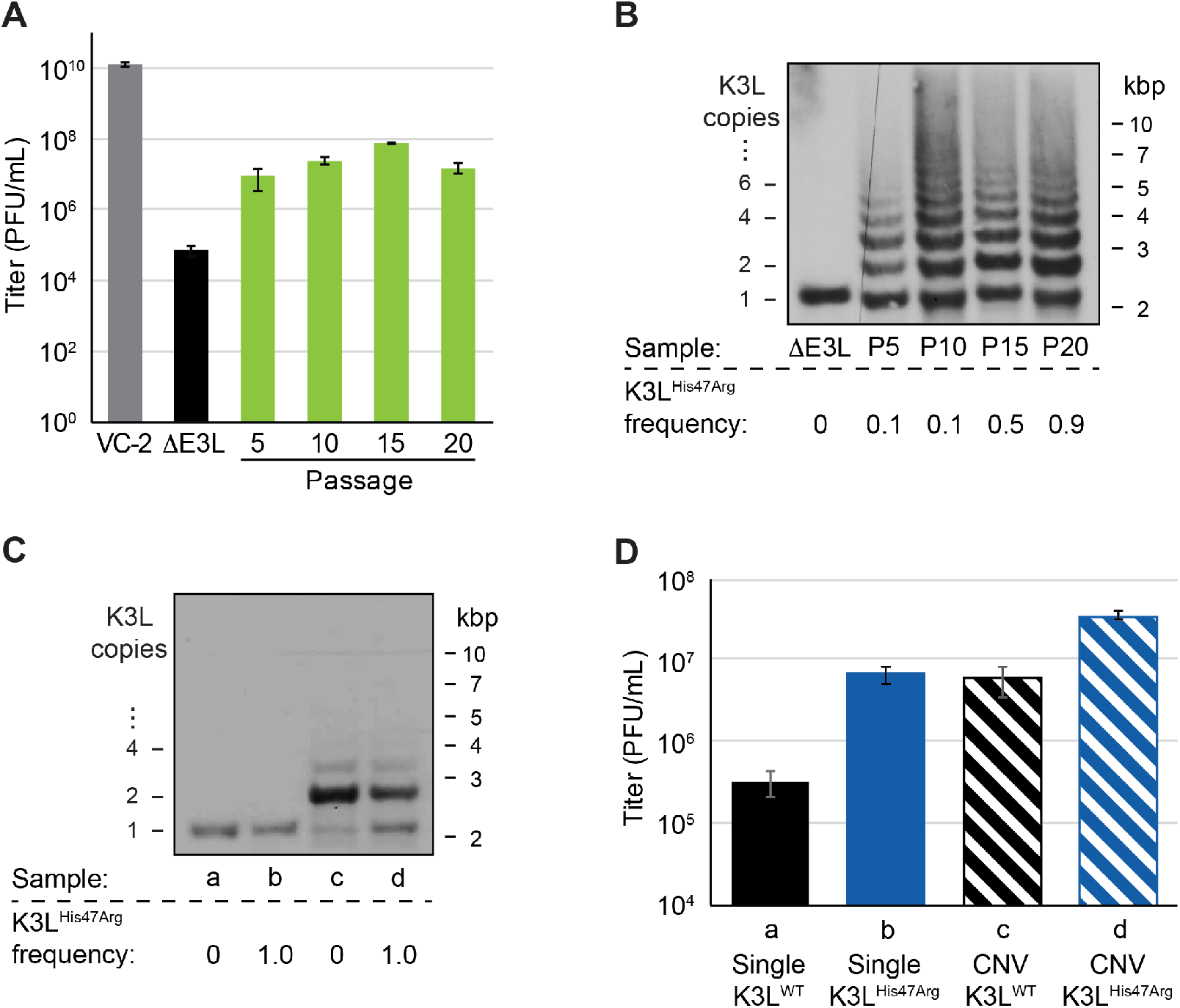
A single nucleotide variant accumulates following increases in K3L copy number. (A) Following 20 serial infections of the ΔE3L strain (MOI 0.1 for 48 hours) in HeLa cells (see **Materials and Methods** for further details), replication was measured in triplicate in HeLa cells for every 5th passage, and compared to wild-type (VC-2) or parent (ΔE3L) virus. (B, C) Digested viral DNA for every 5th passage (B) and four plaque-purified clones (C) was probed with a K3L-specific probe by Southern blot analysis. Number of K3L copies (left) and size in kbp (right) are shown. K3L^His47Arg^ allele frequency for each population (shown below) was estimated by PCR and Sanger sequencing of viral DNA. (D) Replication of plaque purified clones from (C) was measured in HeLa cells in triplicate. All titers were measured in BHK cells by plaque assay, as mean PFU/mL ± standard deviation.

To determine how changes in K3L copy number and the K3L^His47Arg^ mutation might individually contribute to fitness in heterogeneous populations, we isolated distinct variants from single virus clones. Following plaque purification, we obtained viruses containing a single copy of the K3L gene, with or without the K3L^His47Arg^ variant (**Figure 1C**). We also obtained plaque purified clones containing K3L CNV homogenous for either wild-type K3L or K3L^His47Arg^. While viruses with CNV were not clonal due to recurrent recombination between multicopy genomes, they were nearly uniform, harboring mainly 2 copies of K3L following plaque purification (ranging from 1–5; **Figure 1C**), consistent with observations from our previous study (Elde et al., 2012). Comparing the ability of these four viruses to replicate in human cells revealed that either the K3L^His47Arg^ variant or K3L CNV is sufficient for a fitness gain, but the combination of the two genetic changes increases fitness more than either one alone (**Figure 1D**). These results are consistent with earlier reports showing that K3L containing the His47Arg variant is a more potent inhibitor of human PKR than wild-type K3L (Kawagishi-Kobayashi et al., 1997), and that overexpression of the K3L protein is necessary and sufficient to increase viral fitness in human cells (Elde et al., 2012). However, the K3L^His47Arg^ variant only reached high frequency in the population following increases in K3L copy number, suggesting that selection for the variant was affected by copy number. Further tracking the evolution of the virus population thus revealed continued adaptation through genetic changes at the K3L locus.

### Single nucleotide variant accumulation in evolving viral populations

To track the rise of the K3L^His47Arg^ variant in virus populations containing K3L CNV, we needed a means to analyze large and repetitive arrays of sequence. Duplication of K3L results in breakpoints between flanking genetic regions which mark the boundaries of recombination (**Figure 2A**; Elde et al., 2012; Cone et al., 2017). We previously identified two distinct breakpoints flanking K3L in the P10 population that differ by only 3 bp (**Table S1**; Elde et al., 2012), each of which demarcates a duplicon of approximately 500 bp in length (**Figure 2A**). In heterogeneous virus populations with this size of duplicon, short reads cannot discriminate the location of the K3L^His47Arg^ variant either in single copy genomes or within multicopy arrays of K3L (**Figure 2A**). Therefore, we sequenced viral genomes using the ONT MinION sequencing platform, routinely generating reads with a mean length of ∼3 kbp and an N50 between 5–8 kbp (**Table 1**, **Table S2**). These reads, reaching a maximum aligned length of up to 40 kbp, allowed us to directly measure both K3L copy number and the presence or absence of K3L^His47Arg^ in each K3L copy within individual viral genomes (**Figure 2A**). Importantly, we found that differences in flow cell chemistry (and by extension, sequencing error) did not affect our observed distributions of K3L copy number and K3L^His47Arg^ accumulation (**Figure S1**).

**Figure 2.**
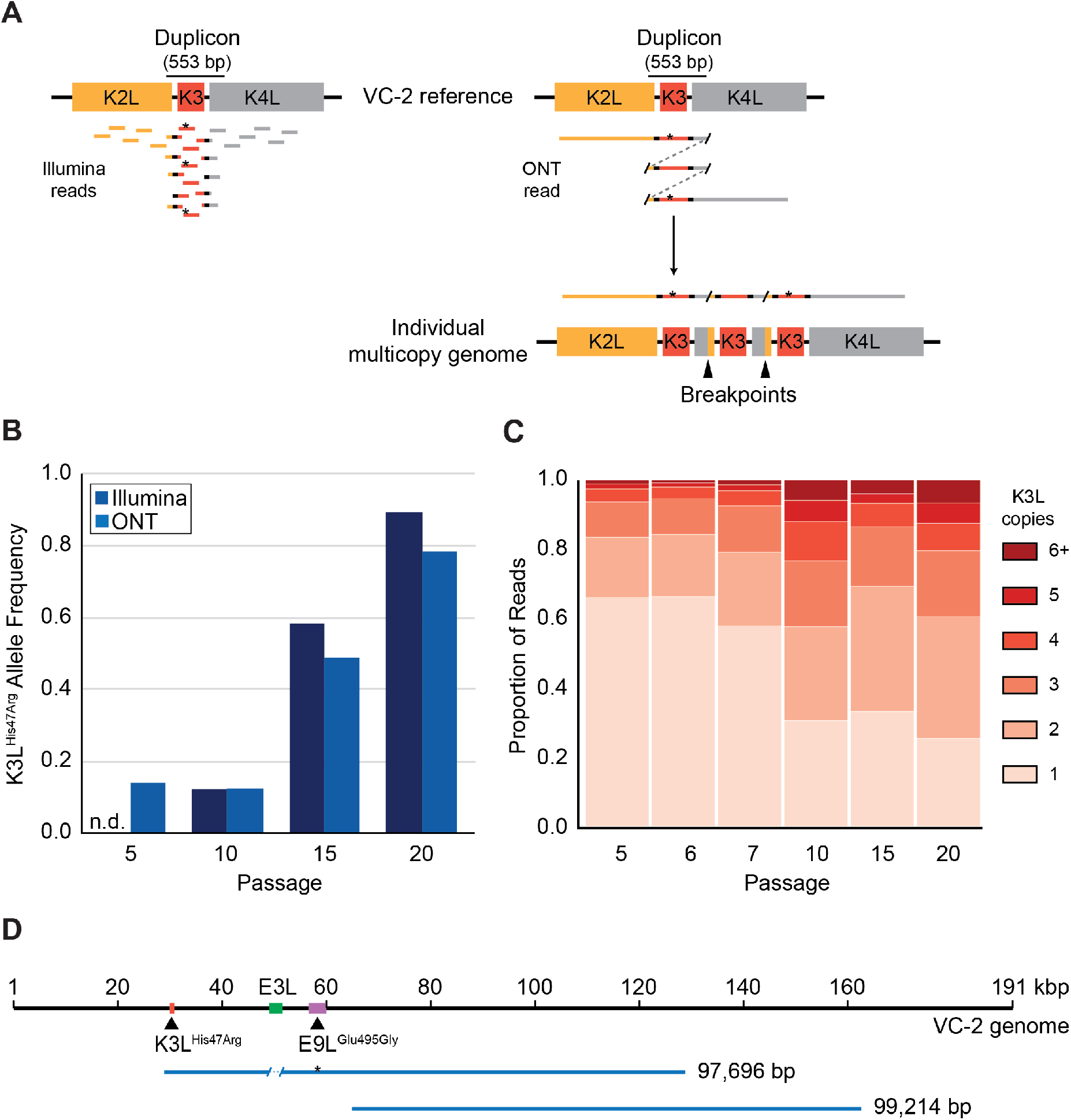
ONT reads capture SNVs and copy number expansions in individual viral genomes. (A) Representative genome structure of the K3L locus in the VC-2 reference is shown on top, with representative Illumina MiSeq and ONT MinION reads shown to scale below. The K3L^His47Arg^ variant within reads is indicated by an asterisk. ONT reads that split and re-align to the K3L duplicon are indicative of individual multicopy genomes (shown below). Tandem duplication breakpoints are indicated by arrowheads. (B) Population-level K3L^His47Arg^ allele frequency was estimated using Illumina or ONT reads from different passages. (C) For each sequenced passage, K3L copy number was assessed within each ONT read that aligned at least once to the K3L duplicon (see **Materials and Methods** for further details). (D) Representative reads from the specific long read preparation are depicted relative to the VC-2 reference genome. The location of the high frequency variants in K3L and E9L (colored blocks) are indicated by arrowheads.

**Table 1.** Summary of ONT sequencing datasets

| Population | Total sequenced reads | Mean read length (bp) | Read length N50 (bp) | Total sequenced bases (Gbp) | Reads containing K3L |
| --- | --- | --- | --- | --- | --- |
| P5 | 239,737 | 2,168 | 5,932 | 0.52 | 1,190 |
| P10 | 91,815 | 3,523 | 7,693 | 0.32 | 914 |
| P15 | 156,829 | 4,249 | 7,067 | 0.67 | 1,599 |
| P20 | 94,050 | 2,893 | 7,702 | 0.27 | 797 |

Analysis of ONT reads from every fifth passage revealed the same two breakpoints previously identified in the P10 population (Elde et al., 2012), suggesting that the 500 bp duplicons were maintained in the viral population over further passages (**Table S1**). Despite a relatively high sequencing error rate (5–10%), variant calling on high frequency (> 0.01) variants using ONT reads yielded similar population-level allele frequencies to those estimated using Illumina MiSeq data for the same samples (**Figure 2B**, **Table S3**; Elde et al., 2012). These results confirm the rapid accumulation of the K3L^His47Arg^ variant between passages 10 and 20, and validate the quality of ONT reads for both structural variant and SNV calling from individual viral genomes.

To assess the full complement of genetic variation in adapting vaccinia populations, we also analyzed ONT data from the different passages to determine if additional mutations had arisen during the course of serial infection. Apart from the K3L^His47Arg^ variant, only one other SNV was identified above an allele frequency of 0.01 in any of the sequenced populationscompared to the parent ΔE3L virus. This variant, a point mutation causing a Glu495Gly amino acid change in the viral DNA polymerase (E9L^Glu495Gly^), decreased in frequency from P10-P20 (**Table S3**), suggesting that it may be non-adaptive. Indeed, while the E9L^Glu495Gly^ SNV reached high frequency at P10 (0.64), this variant alone did not provide a measurable fitness benefit and might have instead accumulated as a hitchhiker mutation (**Figure S2**). These observations, and the lack of any other detectable genetic changes in either the Illumina or ONT data sets, suggest that virus gains in replication were dominated by changes in K3L.

### ONT reads reveal precise K3L copy number in individual genomes

To thoroughly analyze K3L gene amplification in virus populations over time, we first determined K3L copy number within each ONT read from virus populations throughout the passaging experiment (see **Materials and Methods** for further details). Consistent with Southern blot analyses (**Figure 1B**), K3L copy number expansions occurred as early as P5 (**Figure 2C**). Over the course of the next five passages, K3L copy number steadily increased to nearly 70% of virus genomes containing multiple copies of the gene by P10 (**Figure 2C**). From passages 10 to 20, there was a modest shift towards higher copy number genomes, but the distribution of K3L copy number within the population appears to have reached a point near equilibrium. Throughout the experiment, over 90% of viral genomes contained between 1 and 5 copies of K3L (**Figure 2C**), consistent with a fitness trade-off between additional K3L production and increased genome size at very high copy numbers (Elde et al., 2012). However, using ONT, we were able to capture reads from genomes containing up to 15 total copies of K3L (**Figure S3**), suggesting that rare large gene arrays exist in the population but may not persist. These reads reflect the possibility for substantial increases in genome size in adapting virus populations, and highlight the ability of long read sequencing to analyze large arrays of gene repeats.

To test if we had captured a representative sample of large multicopy genomes, we re-sequenced virus genomes from P15 using a DNA library preparation protocol designed to generate extremely long reads (see **Materials and Methods** for further details). High K3L copy number reads could be underrepresented in our initial data sets, because our average sequencing read length of ∼3 kbp may have limited our discovery of viral genomes with greater than 6–8 K3L copies. The alternate protocol produced a mean read length of 9,392 bp (N50 = 19,288 bp), with a maximum aligned read length of 99,214 bp (**Figure 2D**, **Table S2**). Even with increased read lengths, we did not recover larger proportions of high copy vaccinia genomes in this dataset, suggesting that standard library preparation captured a representative sample of K3L copy number in virus genomes. Using this specific long read preparation, we were also able to identify nearly 100 reads that span a ∼30 kbp region separating the two high frequency single nucleotide variants in this population, K3L^His47Arg^ and E9L^Glu495Gly^ (**Figure 2D**). These extremely long sequencing reads enable the direct phasing of distant variants within single poxvirus genomes, and suggest that single reads may routinely capture entire poxvirus genomes in future studies.

### The K3L^His47Arg^ variant rapidly homogenizes in multicopy vaccinia genomes

To determine how single nucleotide variants spread through tandem gene duplications within virus genomes, we scored the presence or absence of the K3L^His47Arg^ variant in each copy of K3L within our ONT reads (see **Materials and Methods** for further details). At P5, the K3L^His47Arg^ SNV was observed almost exclusively in single-copy reads (4 multicopy reads contained the SNV, compared to 304 single-copy reads; **Figure 3**), suggesting that the variant likely originated in a single-copy genome. We further categorized multicopy vaccinia genomes as containing either homogenous K3L arrays (in which every K3L copy contains either the variant or wild-type sequence) or “mixed” arrays (in which both K3L^WT^ and K3L^His47Arg^ copies are present in a single read). In passages 5–7, we observed both mixed and homogenous K3L^His47Arg^ arrays, although mixed arrays were more prevalent (**Figure 3**). The abundance and variety of mixed arrays, combined with the low point mutation rate of poxviruses (Gago et al., 2009; Sanjuán et al., 2010), are consistent with rampant recombination spreading the variant into multicopy genomes, rather than repeated *de novo* mutation or copy number expansion from a single-copy genome containing the K3L^His47Arg^ variant.

**Figure 3.**
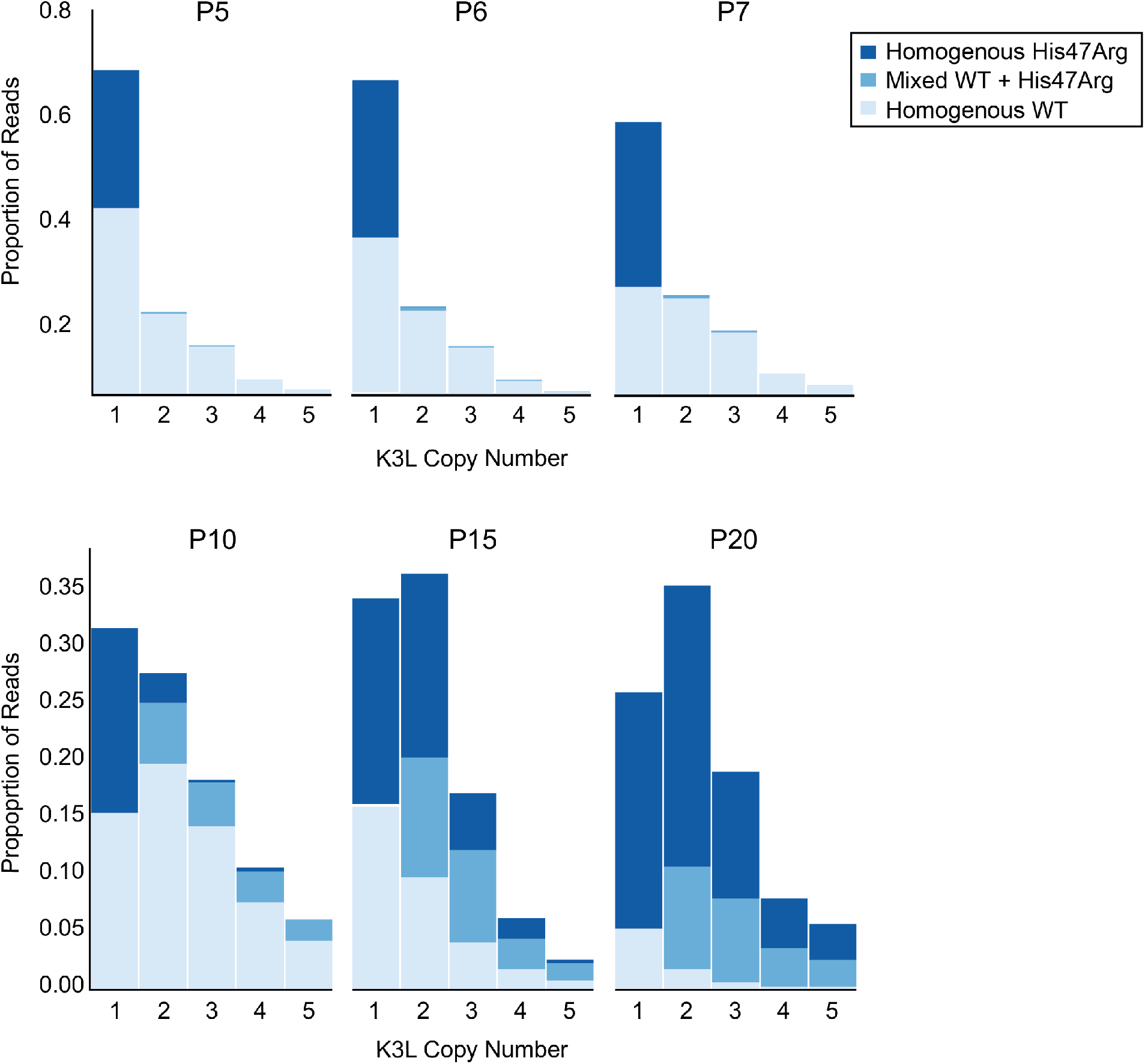
K3L^His47Arg^ variant dynamics within multicopy arrays throughout experimental evolution. Stacked bar plots representing the diversity of allele combinations within single-copy and multicopy reads were generated from ONT reads for the indicated virus populations (passages are listed above each plot). The proportion of reads containing homogenous K3L^WT^, homogenous K3L^His47Arg^, or any combination of mixed alleles is shown for reads containing 1–5 K3L copies.

By P10 of the experiment, when the K3L^His47Arg^ variant was present at a nearly identical population frequency as P5 (0.12 and 0.15, respectively; **Table S3**), the SNV was distributed throughout both single-copy and multicopy reads, although still biased towards single-copy reads (**Figure 3**; **Figure 4**). Strikingly, as K3L^His47Arg^ allele frequency increased in the population by P15 and P20, the SNV was nearly equally distributed throughout genomes at a frequency matching the population-level variant frequency, regardless of K3L copy number (**Figure 4**). These results suggest that once the K3L^His47Arg^ variant entered multicopy genomes, variant accumulation was independent of copy number, and the SNV rapidly became homogenized in gene arrays. Indeed, by P15 and P20, homogenous K3L arrays became increasingly prevalent, and were observed far more than mixed arrays (**Figure 3**). In contrast, simulations of SNV accumulation under a binomial distribution at similar effective population sizes as our experiments (500,000 viruses/serial infection) returned a markedly lower prevalence of homogeneous K3L^His47Arg^ multicopy arrays (**Figure S4**). These findings are consistent with a mechanism of genetic homogenization resembling gene conversion, facilitating the rapid fixation of adaptive variants in homologous sequences.

**Figure 4.**
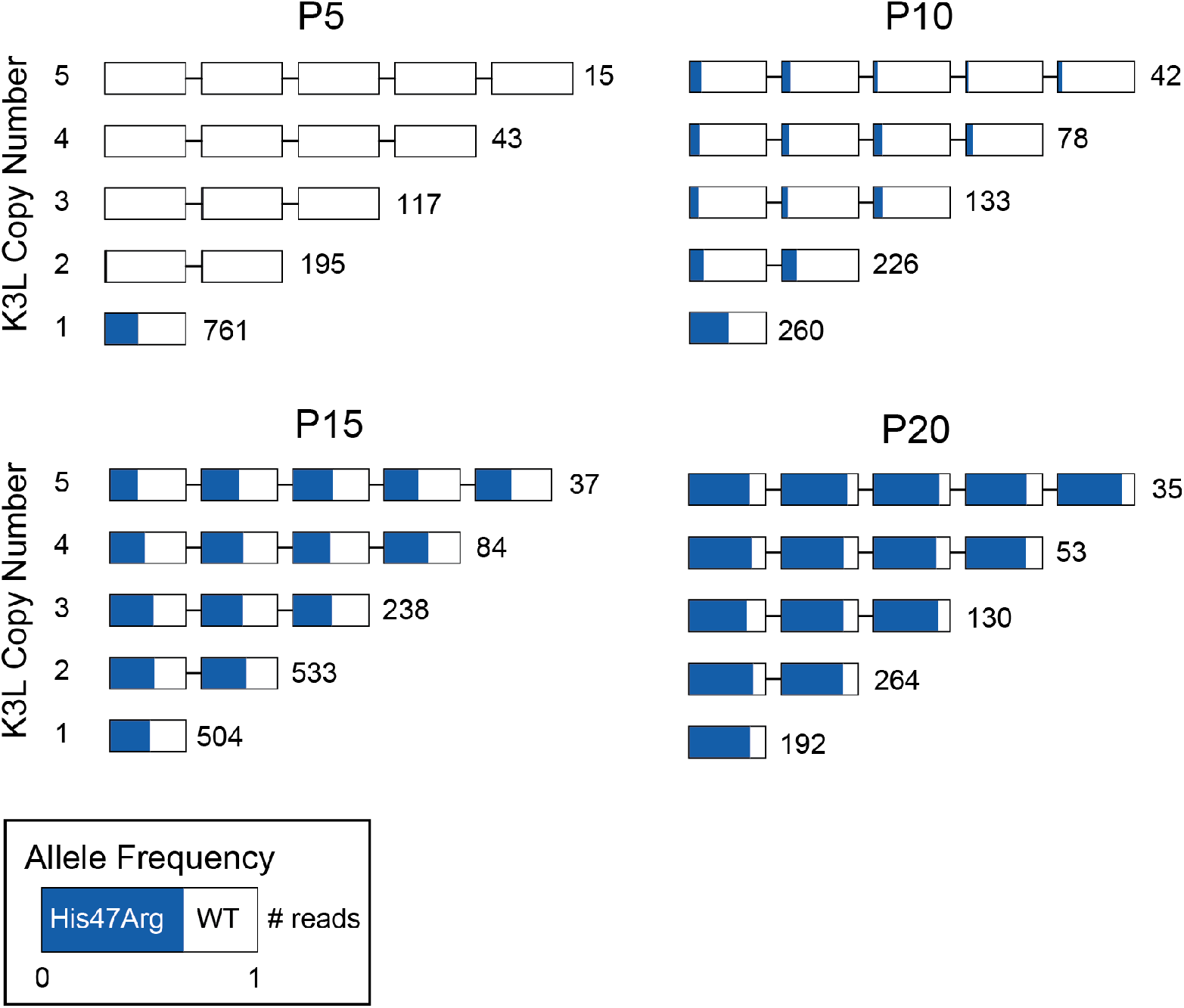
The K3L^His47Arg^ variant rapidly homogenizes in viral genomes regardless of copy number. ONT reads from every 5th passage were grouped by K3L copy number, and each K3L copy was assessed for the presence or absence of the K3L^His47Arg^ SNV. Reads containing 1–5 K3L copies are shown, with the number of reads per group indicated to the right of each row. Reads are oriented 5’ to 3’ relative to the VC-2 reference sequence, and the K3L^His47Arg^ allele frequency in each copy is indicated in blue.

### Recombination and selection drive patterns of K3L^His47Arg^ homogenization

To investigate the influence of intergenomic recombination between co-infecting viruses on gene homogenization, we conducted serial infections at various multiplicity of infection (MOI). We repeated passages 11 to 15 using a range of MOI from 1.0–0.001 (the original experiments are MOI = 0.1), in order to determine whether increasing or virtually eliminating the occurrence of intergenomic recombination would affect the accumulation of the K3L^His47Arg^variant. Analysis of all four P15 populations returned similar distributions of K3L copy number, as well as distributions of homogenous and mixed genomes, regardless of MOI (**Figure 5**). These results suggest that patterns of variant accumulation are robust to a range in MOI of at least four orders of magnitude at each passage. Therefore, intragenomic recombination during replication from single virus infections is likely the main source for rapid homogenization of the K3L^His47A^rg variant within gene arrays. The initial presence of multicopy genomes containing mixed alleles, quickly followed by the predominance of multicopy genomes homogenous for the variant (**Figure 3**), is consistent with abundant intragenomic recombination resulting in gene conversion.

**Figure 5.**
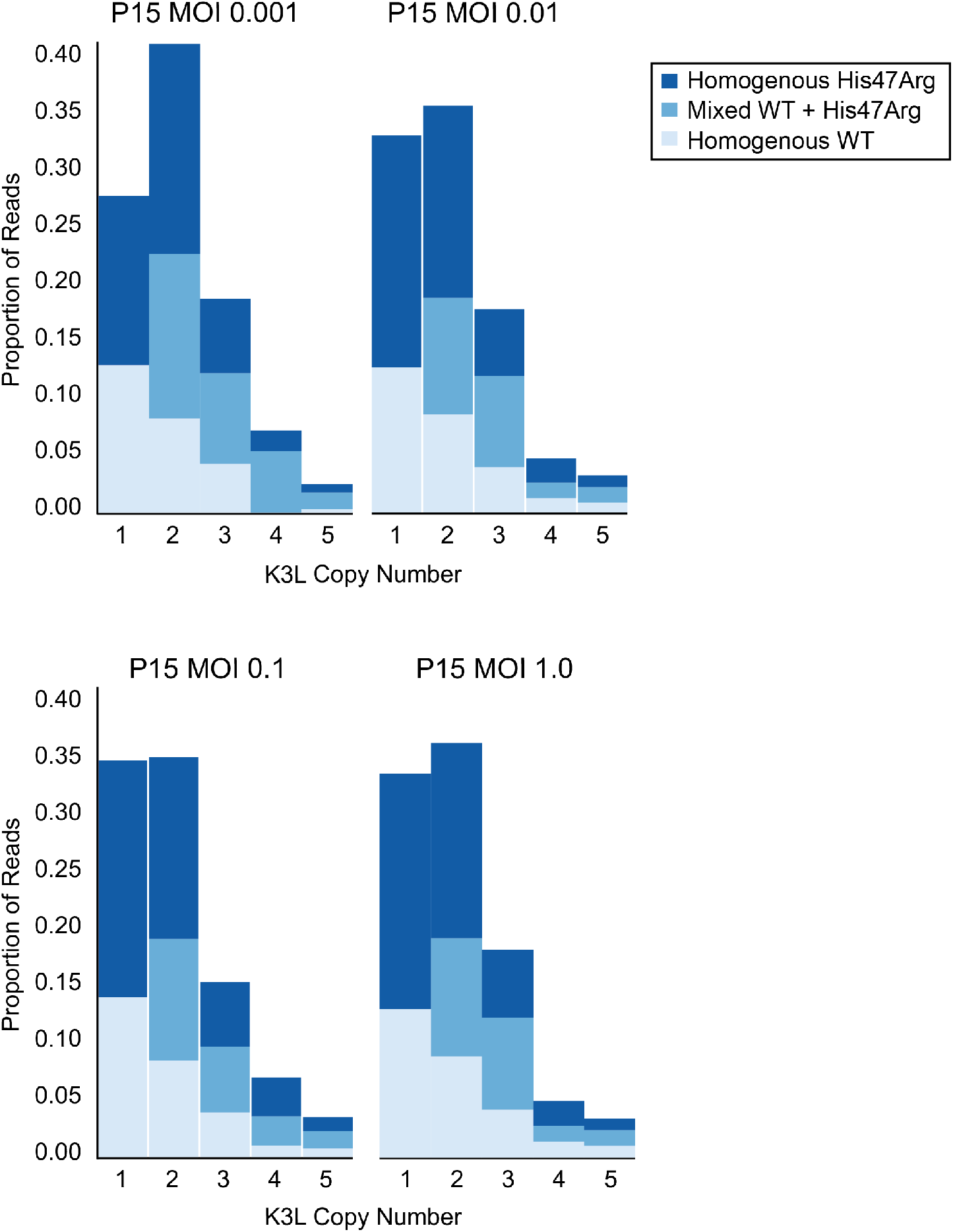
K3L^His47Arg^ homogenization within multicopy genomes is independent of intergenomic recombination rate. The P10 population was serially passaged in HeLa cells at different MOIs (listed above each plot), and each of the resulting P15 populations was sequenced with ONT. Stacked bar plots representing the diversity of allele combinations within single-copy and multicopy reads were generated as in Figure 3.

Gene conversion is a driving force behind sustained homology among repeated sequences, promoting concerted evolution (Chen et al., 2007; Ohta et al., 2010). While the precise role and outcomes of recombination in the spread of the K3L^His47Arg^ variant are difficult to test without a clear understanding of the recombination machinery in poxviruses (Gammon and Evans, 2009), two lines of evidence highlight the importance of natural selection for homogenization of the K3L^His47Arg^ variant in multicopy gene arrays. First, we repeated passages 11 to 15 under relaxed selection on K3L by infecting BHK cells, in which neither the K3L^His47Arg^ variant nor K3L CNV provides a measurable fitness benefit (**Figure S5**; Elde et al., 2012). Consistent with related protocols of plaque purification in BHK cells (**Figure 1C**), we observed a uniform reduction of K3L copy number in the population following 5 passages in BHK cells (P15-BHK) (**Figure 6A**). Further analysis of virus genomes from the P15-BHK population revealed that the K3L^His47Arg^ variant had not increased in frequency compared to P10, in contrast to its rapid accumulation in HeLa cells (**Figure 6B**). Additionally, the proportion of reads homogenous for K3L^His47Arg^ decreased following passaging in BHK cells (**Figure 6C**), as opposed to the rapid homogenization observed during passaging in HeLa cells. These results suggest that variant accumulation and homogenization are dependent on selective pressure imposed by the environment.

**Figure 6.**
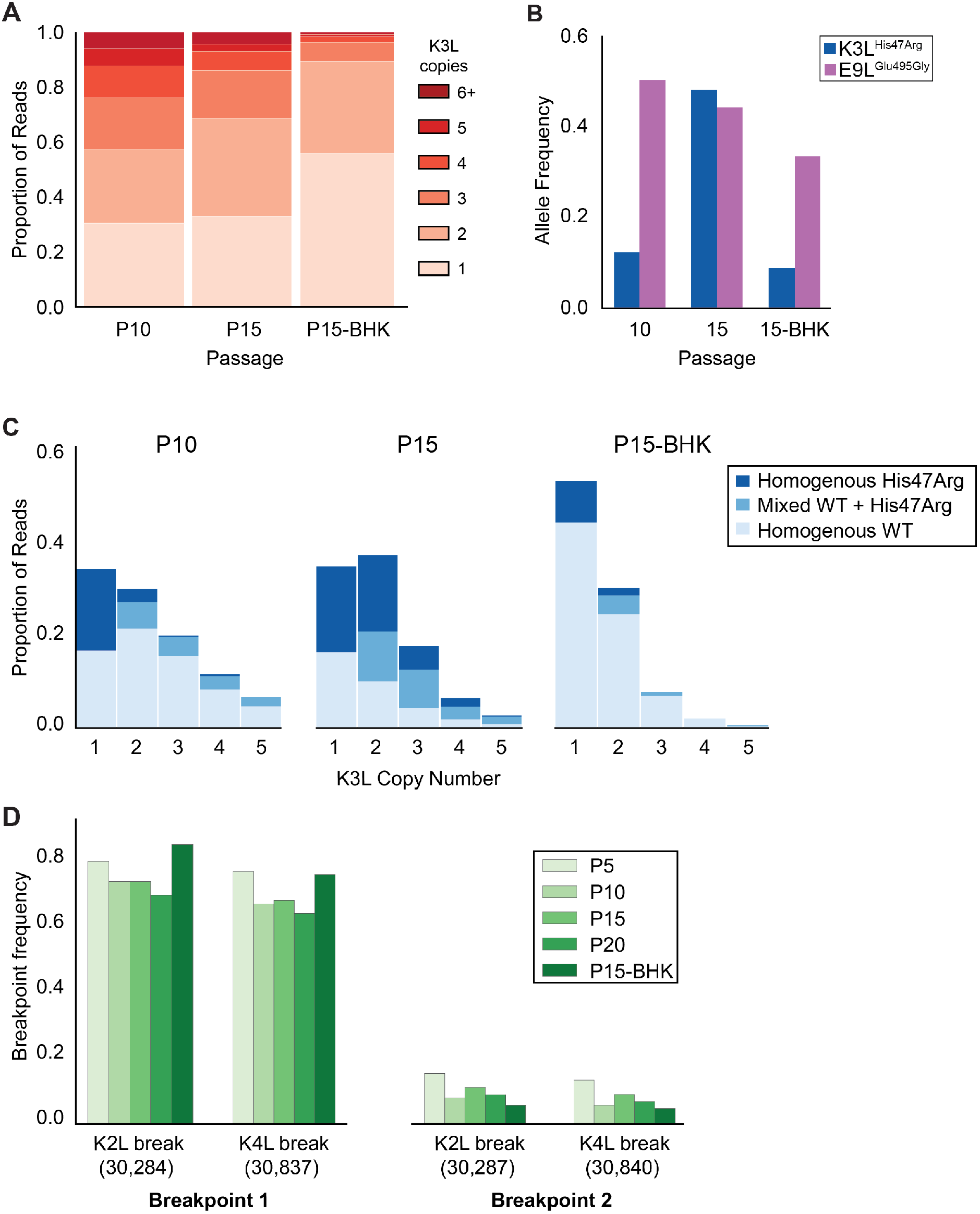
K3L^His47Arg^ variant homogenization is dependent on selection. P10 and P15 data are included from previous figures for comparison with P15-BHK. (A) K3L copy number was assessed for all sequenced reads that unambiguously aligned to K3L at least once, as in **Figure 2C**. (B) K3L^His47Arg^ and E9L^Glu495Gly^ allele frequencies in each population were estimated using ONT reads, as in **Figure 2B**. (C) Stacked bar plots representing the diversity of allele combinations within single-copy and multicopy reads were generated from sequenced ONT reads, as in **Figure 3**. (D) ONT reads were assessed for the presence of each breakpoint by aligning reads to a query sequence containing K3L using BLAST, and extracting the starts and ends of individual alignments to the K3L duplicon. Due to sequencing errors, a proportion of breakpoints do not match either breakpoint 1 or breakpoint 2.

We next compared the accumulation of the K3L^His47Arg^ SNV to variation in the population at two other loci. Two distinct recombination breakpoints (located three base pairs apart in our virus populations) allowed us to compare variation expected to be neutral relative to amino acid position 47 in K3L (**Figure 2A**). Across all analyzed populations, we observed that one breakpoint was dominant over the other, and that the frequency of these breakpoints did not appreciably change over the course of passaging (**Figure 6D, Table S2**). Therefore, while selection for K3L differed between the two cell lines in which viruses were passaged, this did not impact breakpoint frequencies, consistent with a model of neutral variation. In contrast to the rapid accumulation of the K3L^His47Arg^ variant, the only other observed point mutation, E9L^Glu495Gly^, decreased in frequency after passaging in either HeLa or BHK cells (**Figure 6B**). Since only the K3L^His47Arg^ SNV afforded a fitness benefit (**Figure 1D**, **Figure S2**), these results are consistent with selection driving the K3L^His47Arg^ variant to near fixation. The dynamics of these other genetic changes sharply contrast the rapid accumulation of the K3L^His47Arg^ variant in response to selection, suggesting that selection is required to drive rapid homogenization of the K3L^His47Arg^ allele. Together, our results support a gene conversion model of adaptation, in which a beneficial variant that enters an amplified gene array can be spread among the remaining copies through recombination and selection (**Figure 7**).

**Figure 7.**
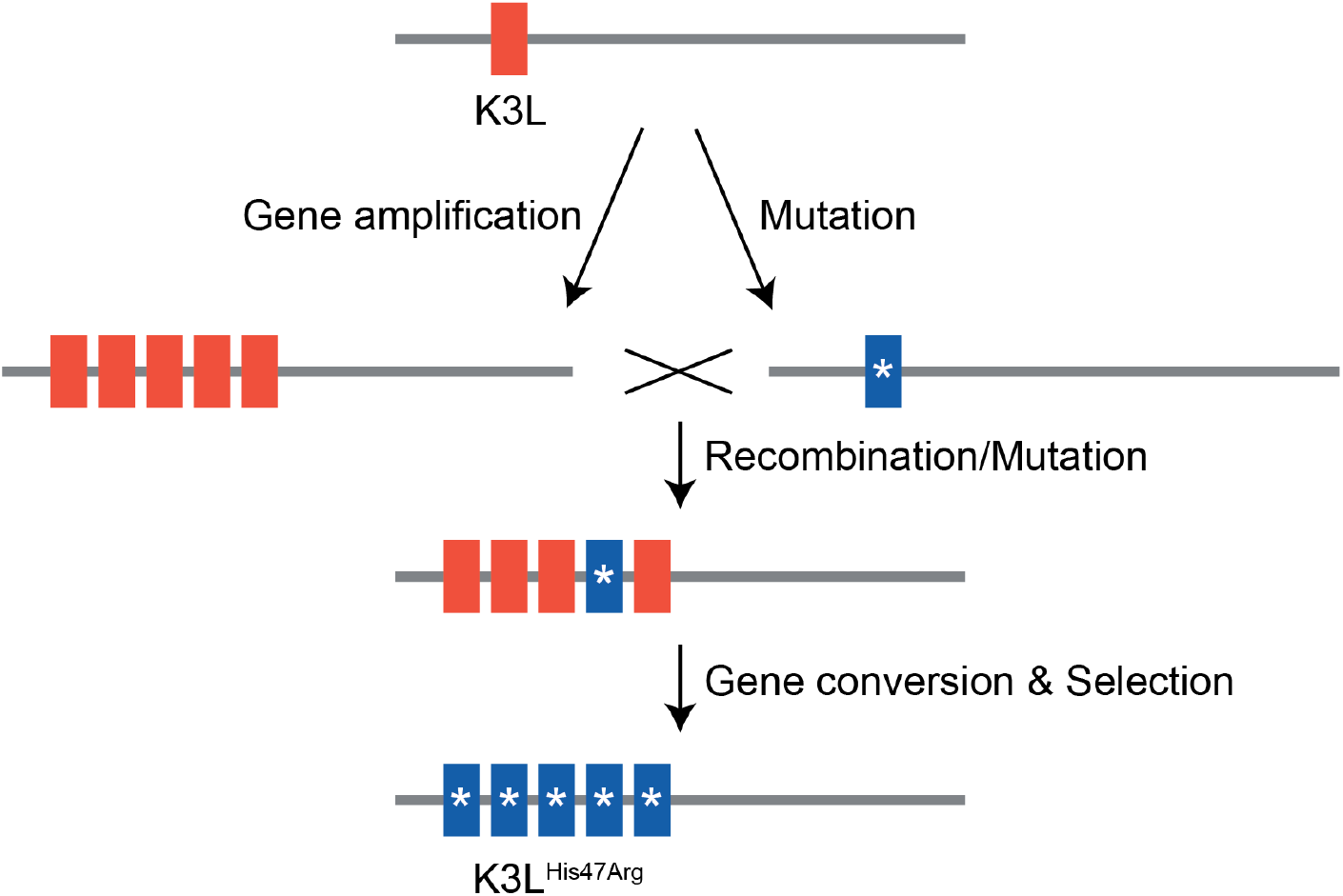
Model of K3L^His47Arg^ homogenization within K3L CNV via gene conversion.

## Discussion

In this study, we investigated how a virus population evolves given the simultaneous presence of distinct adaptive variants at a single locus. In vaccinia virus populations harboring recombination-driven gene copy number amplifications and beneficial point mutations in the same gene, courses of experimental evolution revealed a process of variant homogenization resembling gene conversion. This process could be a unique adaptive feature of large DNA viruses, since evolution through mechanisms of gene duplication are widespread in DNA viruses (McLysaght et al., 2003; Shackelton and Holmes, 2004; Filée 2009; Elde et al., 2012; Filée 2015; Gao et al., 2017), but rare in RNA viruses (Simon-Loriere and Holmes, 2013). For DNA viruses, which possess significantly lower point mutation rates than RNA viruses (Gago et al., 2009; Sanjuán et al., 2010), the rapid fixation of rare beneficial variants within multiple gene copies could be a key to the process of adaptation.

A major outcome from genetic homogenization of beneficial point mutations in multicopy genes might be an enhanced persistence of large gene families. Under this model, the rapid spread of point mutations in gene arrays would counter the advantage of single or low-copy genomes enriched for the SNV to dominate virus populations. Among poxviruses, for example, nearly half of the Canarypox genome consists of 14 gene families, which may have been regularly shaped by mechanisms of genetic homogenization (Afonso et al., 2000; Tulman et al., 2004). Within the emerging classes of giant viruses, the Bodo saltan virus is notable for a large gene family of 148 ankyrin repeat proteins at the ends of its linear genome, with some copies being nearly identical (Deeg et al., 2018). In cases like these, point mutations that are sampled among tandem arrays of genes might quickly spread to fixation through homogenization. Indeed, repeated homogenization through gene conversion has been suggested as the process driving concerted evolution of viral genes (Hughes, 2004), which has been observed in Nanoviruses (Hughes, 2004; Hu et al., 2007; Savory and Ramakrishnan, 2014), Baculoviruses (Zanotto et al., 2008), and Epstein-Barr virus, a human Herpesvirus (Ba abdullah et al., 2017). In these studies, while comparing existing strains or viruses allowed for observations of concerted evolution, short read sequencing of fixed populations restricted the ability to identify an underlying mechanism. In contrast, our experimental system allowed us to track an actively evolving virus population, and using long DNA sequencing reads, uncover a model consistent with gene conversion driving the rapid homogenization of a variant within gene arrays.

Our work also demonstrates the power of long read sequencing to perform high resolution analyses of complex genome dynamics. Using the Oxford Nanopore Technologies platform, we investigated two simultaneous adaptations at single-genome resolution. This type of analysis provides a framework to definitively determine the sequence content of tandem gene duplications, and accurately call variants within these duplicates. Additionally, we demonstrate the ability to phase extremely distant genetic variants, and the longest reads we obtained suggest that entire poxvirus genomes could routinely be captured in single reads. Sequencing entire DNA virus genomes with this level of detail could expand our understanding of DNA virus adaptation as a population evolves, either in an experimental system or during infection of a host. Together, these methods allow for extremely high resolution analyses of complex genomes and could be used to explore the evolution of diverse organisms in new and exciting detail.

## Acknowledgements

We thank Ryan Layer for his assistance in early MinION sequencing experiments, and members of the Quinlan and Elde labs for extremely helpful discussions during preparation of the manuscript. This study was supported by NIH grants R01GM114514 (N.C.E.), R01HG006693 (A.R.Q.), T32AI055434 (K.R.C.), and T32GM007464 (T.A.S.), and an equipment grant from the University of Utah (N.C.E. and A.R.Q.). N.C.E. is a Burroughs Wellcome Fund Investigator in the Pathogenesis of Infectious Disease and H.A. and Edna Benning Presidential Endowed Chair.

## Materials and Methods

### Cells

HeLa and BHK cells were maintained in Dulbecco’s modified Eagle’s medium (DMEM; HyClone, Logan, UT) supplemented with 10% fetal bovine serum (HyClone), 1% penicillin-streptomycin (GE Life Sciences, Chicago, IL), and 1% stable L-glutamine (GE Life Sciences).

### Experimental evolution

The P5-P10 populations of vaccinia virus were previously established following serial passages of the ΔE3L virus (Beattie et al., 1995) in HeLa cells (Elde et al., 2012). Briefly, 150mm dishes were seeded with an aliquot from the same stock of HeLa cells (5×10 ^6^ cells/dish) and infected (MOI = 1.0 for P1, and MOI = 0.1 for subsequent passages) for 48 hours. Cells were then collected, washed, pelleted, and resuspended in 1mL of media. Virus was released by one freeze/thaw cycle followed by sonication. P10 (replicate C passage 10 in Elde et al., 2012) virus was expanded in BHK cells, and titer determined by 48-hour plaque assay in BHK cells performed in triplicate. Passages 11–20 were performed as above, starting with the P10 virus population. Following passage 20, replication ability was assayed simultaneously by 48-hour infection (MOI = 0.1) in triplicate in HeLa cells. For the intergenomic recombination passages, P10 virus was passaged in HeLa cells as above at a range of MOI (MOI = 0.001 - 1.0 as indicated) for 48 hours. For BHK passages, the P10 virus population was passaged five times as above, using an aliquot from the same stock of BHK cells (5×10 ^6^ cells/dish) infected (MOI = 0.1) for 48 hours. All viral titers were determined by 48-hour plaque assay in BHK cells performed in triplicate.

### Southern blot analysis

Viral DNA from purified viral cores was digested with EcoRV (New England Biolabs, Ipswich, MA), and separated by agarose gel electrophoresis. DNA was transferred to nylon membranes (GE Life Sciences) using a vacuum transfer, followed by UV-crosslinking. Blots were probed with PCR-amplified K3L using the DIG High-Prime DNA Labeling & Detection Starter Kit II (Roche, Basel, Switzerland) according to the manufacturer’s protocol.

### Isolation and testing of plaque purified clones

BHK cells were infected for 48 hours with dilutions of P15 or P20 virus, and overlayed with 0.4% agarose. Single plaques were harvested and transferred to new BHK dishes, and resulting wells harvested after 48 hours. Virus was released by one freeze/thaw cycle followed by sonication. The process was repeated three additional times for a total of four plaque purifications. Viral DNA was extracted from individual clones from each population as previously described (Esposito et al., 1981) from infected BHK cells (MOI = 0.1) 24 hours post-infection, and assessed for K3L CNV and the K3L^His47Arg^ SNV by PCR and Sanger sequencing.

One clone of each genotype (single-copy K3L^WT^ or K3L^His47Arg^, and multicopy K3L^WT^ or K3L^His47Arg^) was expanded in BHK cells. Replication ability was assessed by 48-hour infection (MOI = 0.1) in triplicate in either HeLa or BHK cells. Viral titers were determined by 48-hour plaque assay in BHK cells performed in triplicate.

### Deep sequencing of viral genomes

#### Illumina

Total viral genomic DNA was collected as above (Esposito et al., 1981). Libraries were constructed using the Nextera XT DNA sample prep kit (Illumina, Inc., San Diego, CA). Barcoded libraries were pooled and sequenced on an Illumina MiSeq instrument at the High-Throughput Genomics Core (University of Utah, Salt Lake City, UT). Reads were mapped to the Copenhagen reference strain of vaccinia virus (VC-2; accession M35027.1, modified on poxvirus.org; Goebel et al., 1990) using default BWA-MEM v0.7.15 (Li, 2013) parameters as paired-end reads. PCR duplicates were removed using *samblaster* (Faust and Hall, 2014). Variant calling was performed using *freebayes* (Garrison and Marth, 2012), using the following parameters: `freebayes –f $REF $BAM ––pooled-discrete –C 1 ––ploidy 1 ––genotype-qualities ––report-genotype-likelihood-max – F 0.01.`

#### Oxford Nanopore Technologies

Viral particles were isolated from infected BHK cells (MOI = 1.0) 24 hours post-infection, and viral cores were purified by ultracentrifugation through a 36% sucrose cushion at 60,000 rcf for 80 minutes. Total viral genomic DNA was extracted from purified cores as above (Esposito et al., 1981). Purified DNA was then size-selected in a Covaris G-Tube 2 x 1 minute at 6000 rpm. Sequencing libraries for the P10, P15, and P20 populations were prepared using the ONT SQK-NSK007 kit and sequenced on R9 chemistry MinION flow cells (FLO-MIN104); libraries for the P5, P6, and P7 populations were prepared using the ONT SQK-LSK208 kit and sequenced on R9.4 chemistry flow cells (FLO-MIN106); libraries for the P15 MOI 0.01, P15 MOI 0.1, and P15-BHK populations were prepared using the SQK-LSK308 kit and sequenced on R9.5 chemistry flow cells (FLO-MIN107); libraries for the P15 MOI 0.001 and P15 MOI 1.0 populations were sequenced using both R9.4 and R9.5 chemistry flow cells (Oxford Nanopore Technologies Ltd., Oxford, UK). For the specific long read library preparation, we used purified, un-sheared P15 viral DNA and a SQK-RAD002 sequencing kit; libraries were sequenced on a FLO-MIN106 flow cell. All sequencing reactions were performed using a MinION Mk1B device and run for 48 hours; base calling for R9 reactions was performed using the Metrichor cloud suite (v2.40), while a command line implementation of the Albacore base caller (v1.2.4) was used to base call data from the remaining sequencing runs. For Albacore base calling on R9.4 runs, the following command was used: `read_fast5_basecaller.py –k SQK-LSK208 –f FLO-MIN106 –o fast5-t 16 –r –i $RAW_FAST5_DIRECTORY.` For Albacore base calling on R9.5 runs, the following command was used: `full_1dsq_basecaller.py –k SQK-LSK308 –f FLO-MIN107 –o fastq,fast5 –t 16 –r –i $RAW_FAST5_DIRECTORY.` For all R9 and R9.4 data, FASTQ sequences were extracted from base-called FAST5 files using *poretools* (Loman and Quinlan, 2014), while FASTQ were automatically generated by Albacore during base-calling on the R9.5 populations. Prior to alignment, adapter trimming on all ONT reads was performed using Porechop (https://github.com/rrwick/Porechop). Only the highest quality reads (2D and 1D^2^ for R9/R9.4 and R9.5 chemistries, respectively) were used for downstream analysis. Pooled FASTQ files for each sample were then aligned to the VC-2 reference genome with BWA-MEM v0.7.15 (Li, 2013), using the default settings provided by the `–xont2d` flag. Population-level estimates of SNV frequencies were determined from our nanopore data using *nanopolish* v0.8.4 (Loman et al., 2015).

### Copy number and allele frequency estimation

Custom Python scripts (github.com/tomsasani/vacv-ont-manuscript) were used to calculate both K3L copy number and K3L^His47Arg^ allele frequency within individual aligned viral reads. Briefly, to identify individual ONT reads containing K3L, we first selected all reads that aligned at least once to the duplicon containing K3L. We next categorized ONT reads containing K3L as single-copy or multicopy. Reads that unambiguously aligned once to the K3L locus were classified as single-copy. The VC-2 reference contains a single K3L gene; therefore, if more than one distinct portion of a read aligned to the full length of K3L, that read was instead classified as multicopy. We further filtered reads to include only those that aligned to the K3L duplicon, as well as 150 bp of unique VC-2 sequence upstream and downstream of the K3L duplicon, to ensure that the read fully contained the estimated number of K3L copies. Finally, we removed sequencing reads with one or more truncated alignments to the K3L duplicon, or that contained any alignments to K3L with mapping qualities less than 20. All Python code used to determine copy number and identify the K3L^His47Arg^ SNV within K3L arrays is available in the GitHub repository, in addition to instructions for plotting/visualizing the data.

### Breakpoint characterization

To characterize the proportions of K3L duplicon breakpoint pairs in the ONT data, we first extracted all reads from each population that unambiguously aligned to K3L at least once, and that also aligned to unique sequence 150 base pairs up- and downstream of the K3L duplicon. We then aligned a 1000 bp query sequence (containing VC-2 reference sequence from genomic position 30,000 to 31,000) to each of the extracted reads using BLAST (Zhang et al., 2000). For every read, we extracted the start and end coordinates of all high-quality alignments to the K3L query.

### Estimating ONT sequencing error

In order to estimate ONT sequencing error in each population of viral reads, we randomly sampled 1000 sites in the vaccinia VC-2 reference known not to be polymorphic in our passaged virus population. We then calculated the alternate allele frequency at each site by dividing total alternate support in the corresponding BAM (mismatches, insertions, and deletions) by the total number of nucleotides aligned to the site. Alternate allele frequency estimates were then averaged across all 1000 sites to obtain a proxy for sequencing error.

### Generating simulated distributions of the K3L^His47Arg^ variant within multicopy arrays

To visualize a simulated distribution of the K3L^His47Arg^ allele in multicopy K3L genomes, we first created a hypothetical population of 500,000 vaccinia virus genomes for each population, which matched the copy number distribution of the passage of interest. We then selected a genome from this population at random, and selected a random copy within that genome. After randomly sampling a single value from a uniform distribution (0.0 to 1.0, incremented by 0.01), if that value was less than or equal to the observed population allele frequency of K3L^His47Ar^g, we “mutated” the selected copy. This process was repeated until the population allele frequency of the hypothetical population matched the observed population allele frequency at that passage. This process effectively simulated a binomial distribution of K3L^His47Arg^ alleles within the hypothetical population, with probability *p* equal to the observed K3L^His47Arg^ allele frequency in the passage.

### Accession numbers

All deep sequencing data will be available in FASTQ format on the Sequence Read Archive. Previously published Illumina MiSeq reads from the P10 population are available under accession SRP013146 (Elde et al., 2012).

**Table S1:** Structural variant breakpoint frequencies during passaging.

| **Breakpoint** | **K2L break** | **K4L break** | **P5** | **P10** | **P15** | **P20** | **P15-BHK** |
| --- | --- | --- | --- | --- | --- | --- | --- |
| 1 | 30,284 | - | 0.76 | 0.69 | 0.69 | 0.65 | 0.83 |
| 1 | - | 30,837 | 0.76 | 0.63 | 0.64 | 0.61 | 0.75 |
| 2 | 30,287 | - | 0.14 | 0.06 | 0.10 | 0.08 | 0.05 |
| 2 | - | 30,840 | 0.12 | 0.04 | 0.07 | 0.05 | 0.04 |

**Table S2:** Complete summary of ONT sequencing datasets.

| **Population** | **Total sequenced reads** | **Mean read length (bp)** | **Read length N50 (bp)** | **Total sequenced bases (Gbp)** | **Reads containing K3L** |
| --- | --- | --- | --- | --- | --- |
| P5 | 239,737 | 2,168 | 5,932 | 0.52 | 1,190 |
| P6 | 163,991 | 4,530 | 7,738 | 0.74 | 2,125 |
| P7 | 220,230 | 4,007 | 7,051 | 0.88 | 1,645 |
| P10 | 91,815 | 3,523 | 7,693 | 0.32 | 912 |
| P15 | 156,829 | 4,249 | 7,067 | 0.67 | 1,599 |
| P20 | 94,050 | 2,893 | 7,702 | 0.27 | 789 |
| P15 - MOI 0.001 | 55,140 | 2,341 | 5,360 | 0.13 | 315 |
| P15 - MOI 0.01 | 128,774 | 2,417 | 6,078 | 0.31 | 865 |
| P15 - MOI 0.1 | 171,765 | 2,208 | 5,696 | 0.38 | 979 |
| P15 - MOI 1.0 | 67,391 | 2,704 | 4,264 | 0.18 | 365 |
| P15 - BHK | 21,675 | 5,108 | 8,141 | 0.12 | 373 |

**Table S3:**
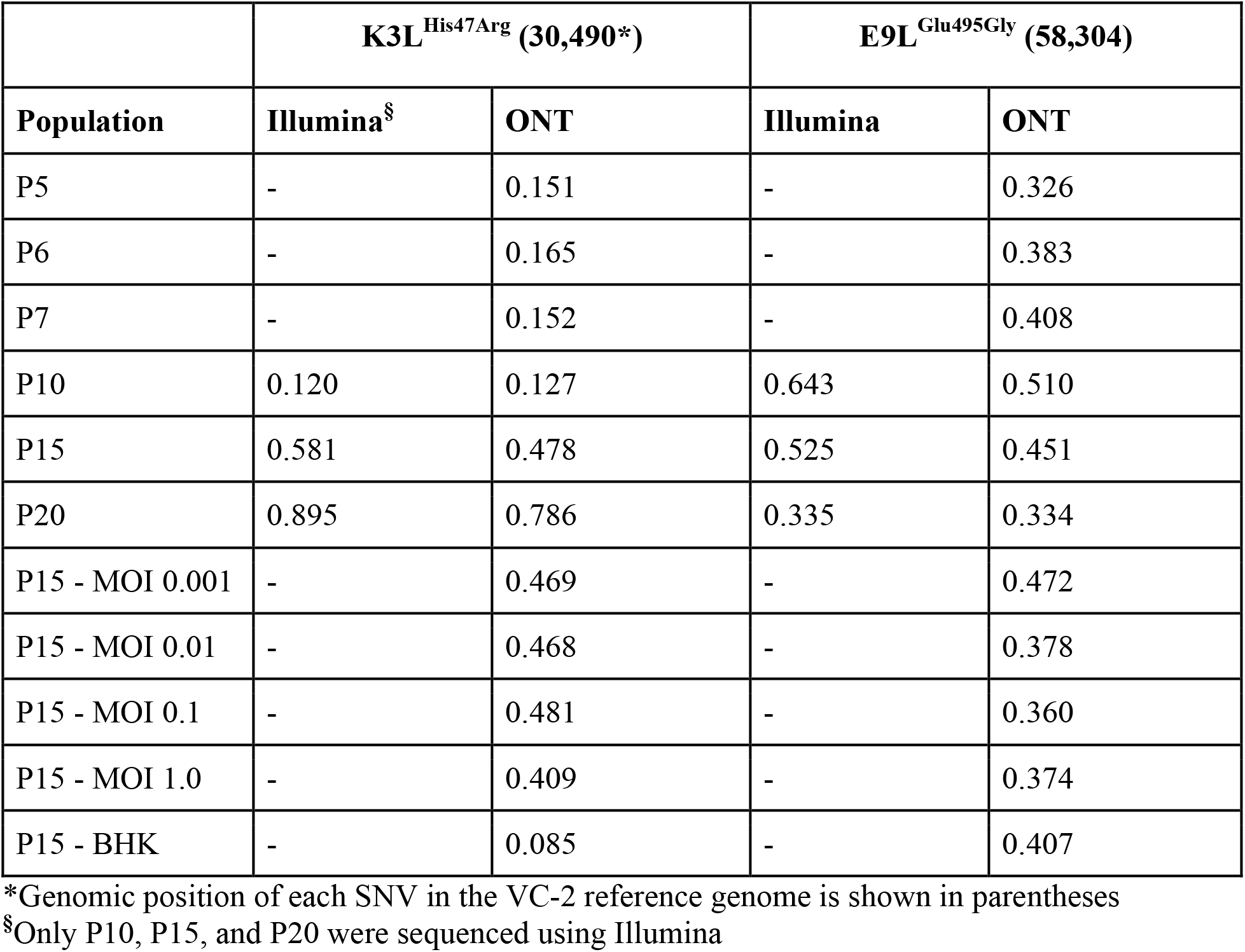
Single nucleotide variants in virus populations from Illumina or ONT datasets.

|  | **K3L<sup>His47Arg</sup> (30,490\*)** |  | **E9L<sup>Glu495Gly</sup> (58,304)** |  |
| --- | --- | --- | --- | --- |
| **Population** | **Illumina<sup>§</sup>** | **ONT** | **Illumina** | **ONT** |
| P5 | - | 0.151 | - | 0.326 |
| P6 | - | 0.165 | - | 0.383 |
| P7 | - | 0.152 | - | 0.408 |
| P10 | 0.120 | 0.127 | 0.643 | 0.510 |
| P15 | 0.581 | 0.478 | 0.525 | 0.451 |
| P20 | 0.895 | 0.786 | 0.335 | 0.334 |
| P15 - MOI 0.001 | - | 0.469 | - | 0.472 |
| P15 - MOI 0.01 | - | 0.468 | - | 0.378 |
| P15 - MOI 0.1 | - | 0.481 | - | 0.360 |
| P15 - MOI 1.0 | - | 0.409 | - | 0.374 |
| P15 - BHK | - | 0.085 | - | 0.407 |

\*Genomic position of each SNV in the VC-2 reference genome is shown in parentheses
§Only P10, P15, and P20 were sequenced using Illumina

**Figure S1.**
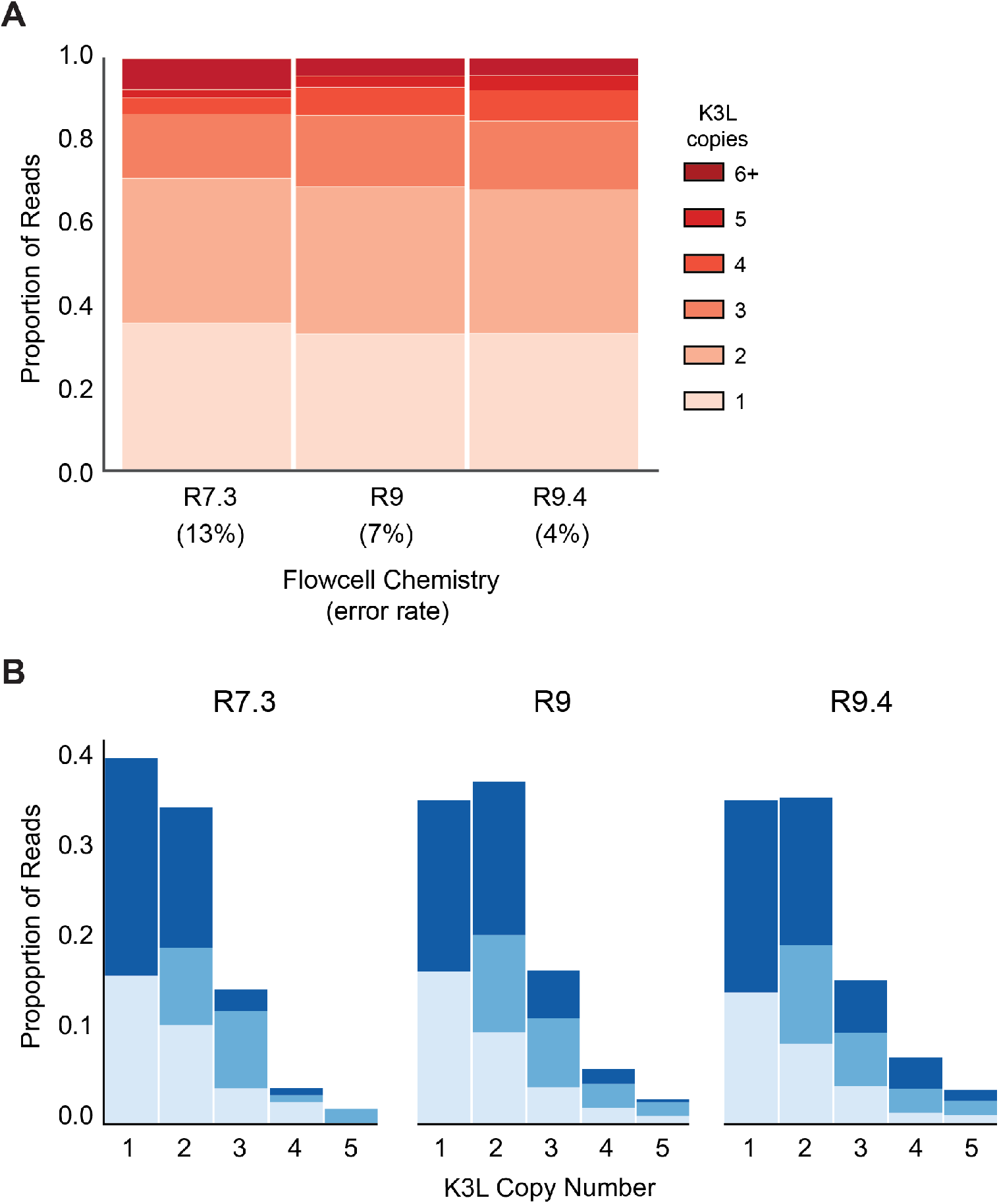
ONT sequencing error rate does not impact inferred patterns of K3L copy number or K3L^His47Arg^ allele frequency. (A) The P15 vaccinia population was sequenced with R7.3, R9, and R9.4 chemistry ONT flowcells, with sequencing error rates of approximately 13%, 7%, and 4%, respectively (see **Materials and Methods** for further detail). K3L copy number was assessed in each sequenced population as described in **Figure 2C**. (B) Stacked bar plots representing the diversity of allele combinations within single-copy and multicopy reads were generated from sequenced ONT reads from each chemistry, as in **Figure 3** and **Figure 5**.

**Figure S2.**
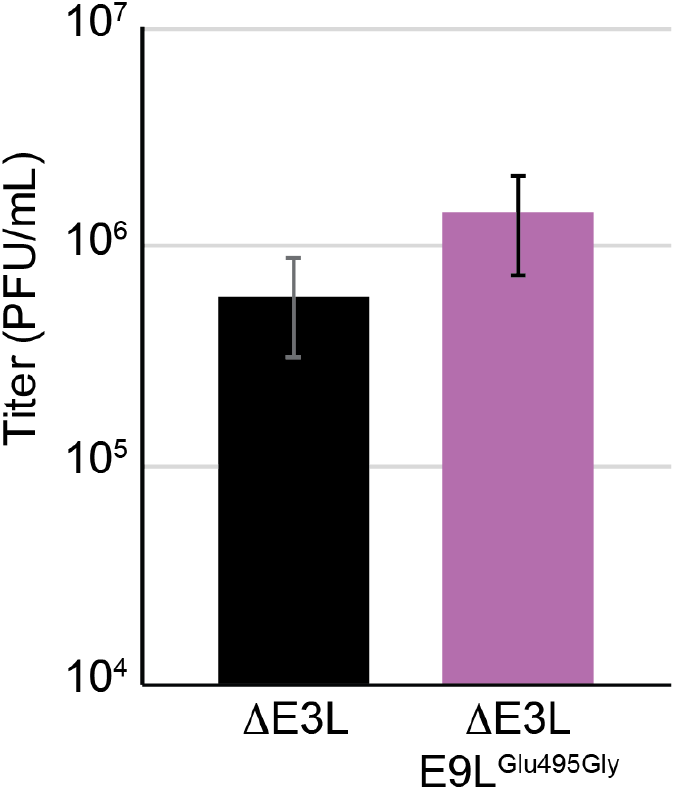
The E9L^Glu495Gly^ variant does not contribute to viral fitness. A virus clone containing the E9L^Glu495Gly^ variant as the only genetic change relative to ΔE3L was identified following four rounds of plaque purification in BHK cells (clone a in **Figure 1C**). Replication was measured compared to ΔE3L virus by 48 hour infection (MOI 0.1) of HeLa cells in triplicate. Titers were measured in BHK cells by plaque assay, as mean PFU/mL ± standard deviation.

**Figure S3.**
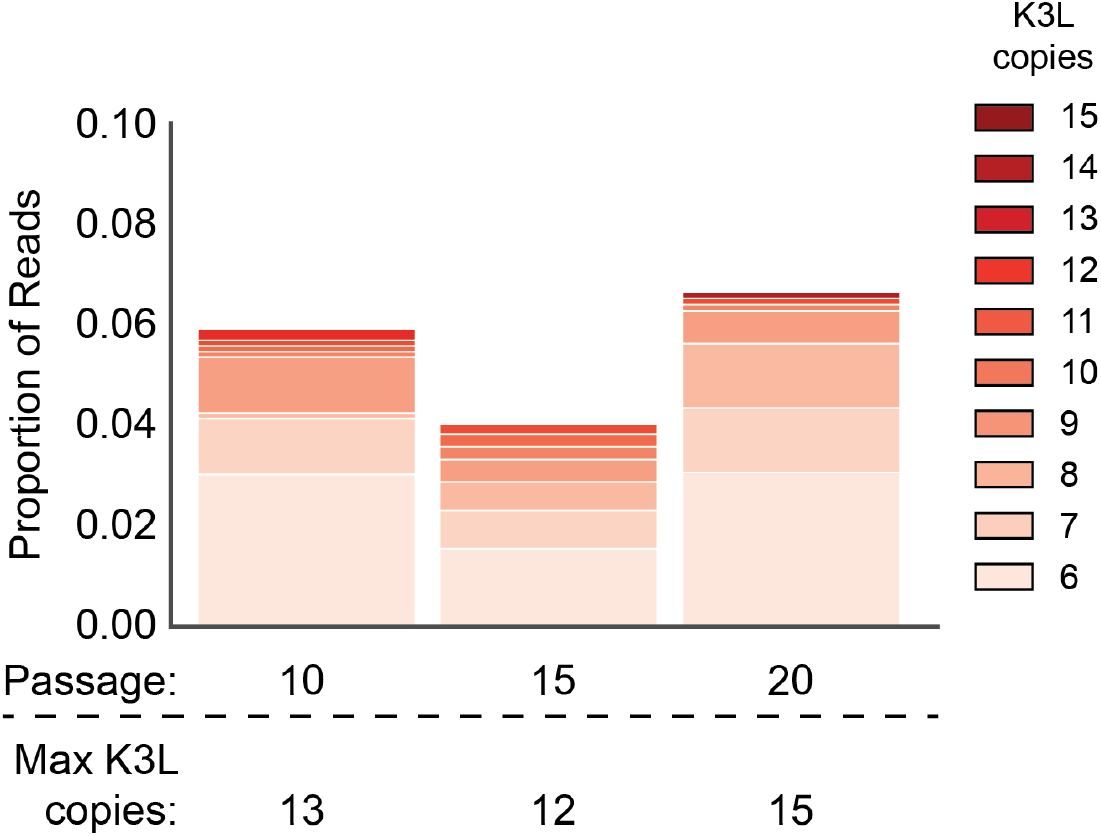
ONT reads capture extremely high K3L copy number in vaccinia genomes. K3L copy number was assessed in ONT reads from P10, P15, and P20 as described in **Figure 2C** Stacked bar plots indicate overall proportions of sequencing reads that contain between 6 and 15 copies of K3L, in ascending order (darker bars represent increasing copy number). The maximum copy number observed in any single ONT read is indicated below each passage.

**Figure S4.**
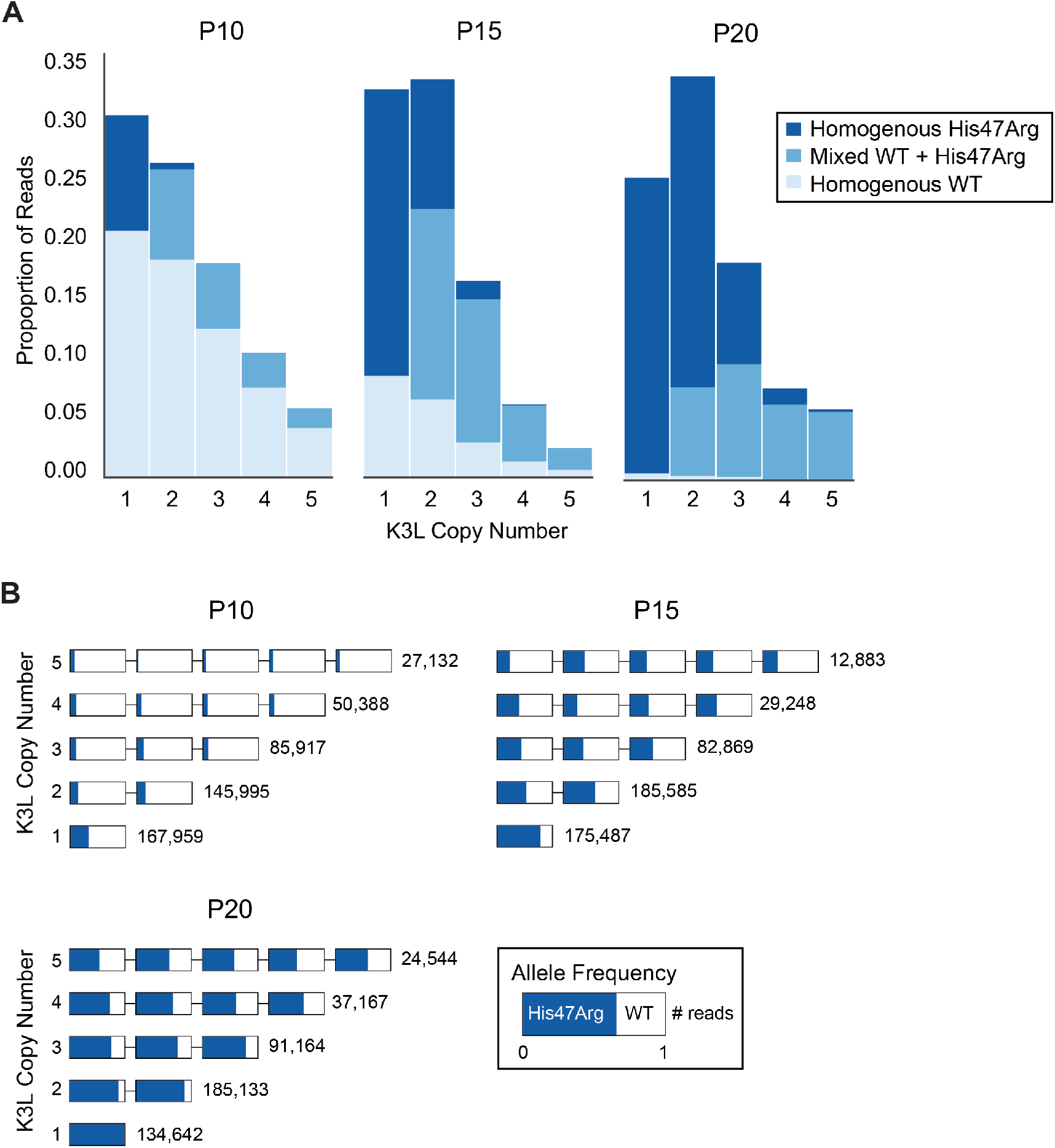
Simulated accumulation of the K3L^His47Arg^ allele. (A) The K3L^His47Arg^ allele was uniformly distributed in simulated vaccinia populations with copy number distributions identical to passages P10, P15, and P20 (see **Materials and Methods** for further detail). Stacked bar plots representing the diversity of allele combinations within single-copy and multicopy genomes were generated as in **Figure 3** and **Figure 5**. (A) The frequency of the K3L^His47Arg^ variant in each copy within genomes of between 1 and 5 copies of K3L for the simulated populations was generated as in **Figure 4**.

**Figure S5.**
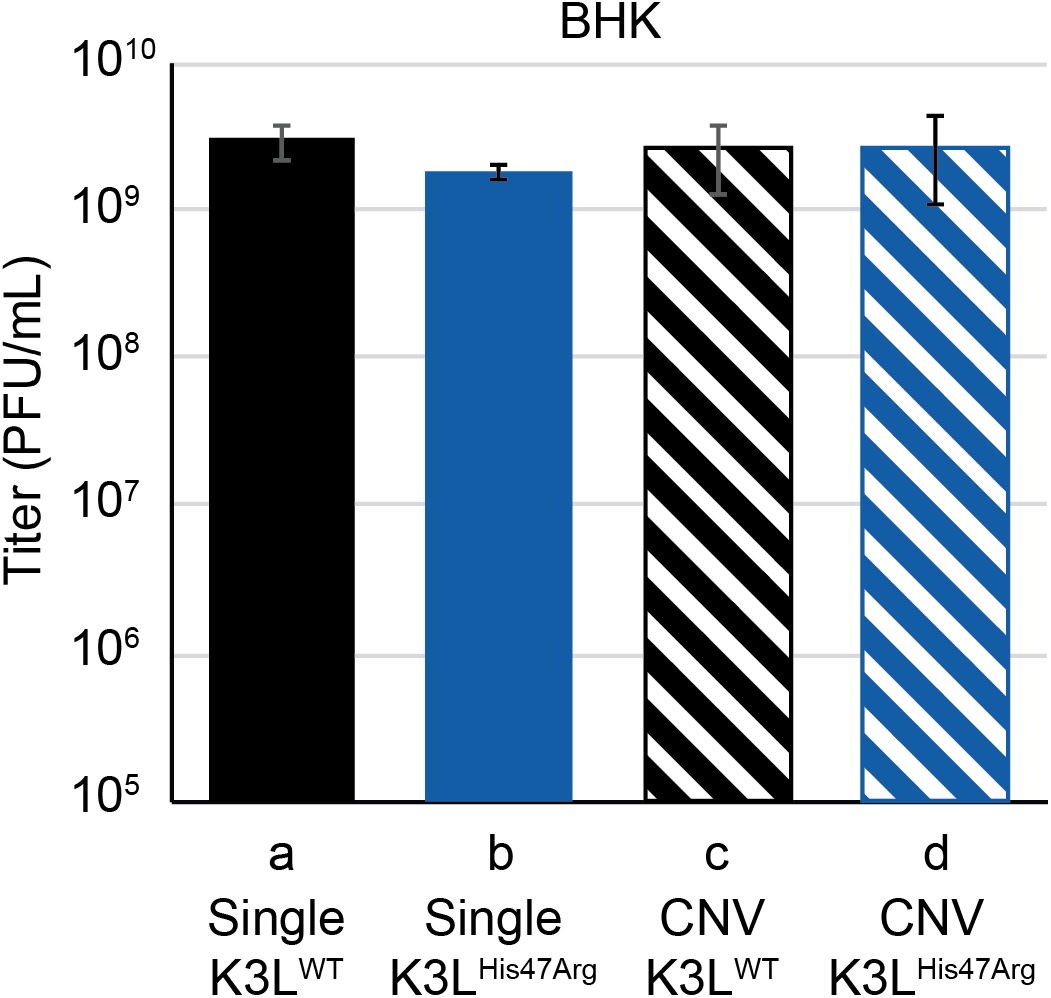
K3L^His47Arg^ and K3L CNV are non-adaptive in the permissive BHK cell line. Replication was measured for plaque purified clones (as in **Figure 2C**) in BHK cells by 48 hour infection (MOI 0.1) in triplicate. All titers were measured in BHK cells by plaque assay, as mean PFU/mL ± standard deviation.

